# *in vivo* expression vector derived from anhydrobiotic tardigrade genome enables live imaging in Eutardigrada

**DOI:** 10.1101/2022.09.21.508853

**Authors:** Sae Tanaka, Kazuhiro Aoki, Kazuharu Arakawa

**Affiliations:** Institute for Advanced Biosciences, Keio University, Japan; Exploratory Research Center on Life and Living Systems (ExCELLS), National Institutes of Natural Sciences, Japan; National Institute for Basic Biology, National Institutes of Natural Sciences, Japan; Faculty of Life Science, SOKENDAI (Graduate University for Advanced Studies), Japan; Graduate School of Media and Governance, Keio University, Japan; Faculty of Environment and Information Studies, Keio University, Japan

**Keywords:** anhydrobiosis, tardigrades, *in vivo* expression, live imaging

## Abstract

Water is essential for life, but anhydrobiotic tardigrades can survive almost complete dehydration. Anhydrobiosis has been a biological enigma for more than a century with respect to how organisms sustain life without water, but the few choices of genetic toolkits available in tardigrade research have been a challenging circumstance. Here, we report the development of an *in vivo* expression system for tardigrades (the TardiVec system). TardiVec is based on a plasmid vector with promoters that originated from an anhydrobiotic tardigrade *Ramazzottius varieornatus*. It enables the introduction of GFP-fused proteins and genetically encoded indicators such as the Ca^2+^ indicator GCaMP into tardigrade cells; consequently, the dynamics of proteins and cells in tardigrades may be observed by fluorescence live imaging. This system is applicable for several tardigrades in the class Eutardigrada: the promoters of anhydrobiosis-related genes showed tissue-specific expression in this work. Surprisingly, promoters functioned similarly between multiple species, even for species with different modes of expression of anhydrobiosis-related genes, such as *Hypsibius exemplaris*, in which these genes are highly induced upon facing desiccation, and *Thulinius ruffoi*, which lacks anhydrobiotic capability. These results suggest that the highly dynamic expression changes in desiccation-induced species are regulated in *trans*. Tissue-specific expression of tardigrade-unique unstructured proteins also suggests differing anhydrobiosis machinery depending on the cell types. We believe that TardiVec opens up various experimental possibilities in tardigrade research, especially to explore anhydrobiosis mechanisms.

## Introduction

Water is essential for life on Earth. In contrast, some organisms, e.g., yeast, artemia and tardigrades, can enter an almost completely dehydrated state, referred to as anhydrobiosis^1^. Anhydrobiotic organisms halt their metabolic activity and maintain the structures of living systems under water deficit conditions until they resume active metabolism upon rehydration. Tardigrades in the anhydrobiotic state can tolerate various extreme conditions, e.g., temperature at almost absolute zero^2^, high pressure (7.5 GPa)^3^, high doses of irradiation of UV and gamma rays^4–11^, and even exposure to space vacuum^12–14^.

In terrestrial tardigrades, the mechanisms underlying anhydrobiosis have been gradually revealed through analyses of genomes and transcripts^15,16^. Tardigrades have evolved lineage-specific IUP (intrinsically unstructured protein) families localized in the cytosol, nucleus, mitochondria and extracellular space, i.e., CAHS (cytoplasmic abundant heat-soluble), Dsup (damage suppressor), MAHS (mitochondrial abundant heat-soluble) and SAHS (secretory abundant heat-soluble)^17–19^. Previous studies used heterologous hosts for molecular analyses of these proteins and showed that exogenous expression of Dsup and MAHS confers tolerance to human cells^18,19^ and plants^20,21^, and a deficit of CAHS transcripts by RNAi attenuated the survival rate in the dehydrated state^22^; however, little is known about the actual tissue specificity or the actual workings of these tardigrade-specific IUPs within tardigrades during anhydrobiosis due to the lack of a genetic toolkit for use in these species.

Recently, several reports indicated that recombinant CAHS proteins form fibers or a gel-like lump at high concentrations *in vitro*, and similar fibrous condensates or aggregates were observed in human cultured cells and E. coli cells^23–27^. These gel-like structures are assumed to sustain the structure of a whole cell or to prevent membranes and proteins from cohesion in the cytoplasm. This concept follows the research of another anhydrobiosis-related IUPs, namely, the LEA (late embryogenesis abundant) protein, one of which from artemia is reported to form liquidlJliquid phase separation (LLPS) upon dehydration *in vitro*^*28*^. Interestingly, such condensation is rapidly reversible; that is, condensates formed with applied osmotic pressure rapidly dissociate upon removal of such stress^23^. This highly dynamic nature demands a live imaging toolkit for *in vivo* study. On the other hand, reports thus far are all based on *in vitro* experiments or heterologous systems, such as human culture cell lines, yeast, and bacteria, since direct approaches applicable in tardigrades are currently restricted to RNAi and IHC (immunohistochemistry). Therefore, the actual subcellular localization in tardigrade cells and the dynamics during anhydrobiosis, as well as the tissue specificity of these expressions in tardigrades, remain elusive. IHC has been used widely in tardigrades to reveal cell morphology, such as muscle and neural systems, and the localization of tardigrade protein in the whole body^29–36^, but it nevertheless cannot be used to observe the dynamics of proteins and cells in a living tardigrade.

In the present study, we have developed an *in vivo* expression system, designated the TardiVec system, using a promoter that originated from an anhydrobiotic tardigrade, *Ramazzottius varieornatus*; it allows the introduction of exogenous genes such as GFP for live imaging in tardigrades. Tardigrade vectors show distinct expression patterns depending on the promoter sequence, and the GFP signals were retained for more than 10 days in tardigrades. The genetically encoded calcium indicator GCaMP^37^ functions in the muscle and neurons of tardigrades, and its function is maintained throughout anhydrobiosis and after rehydration. We also confirmed that promoters that originated from not only housekeeping genes but also tardigrade-specific genes function widely in other tardigrade species within the order Parachela, and to some extent within the class Eutardigrada. Furthermore, the TardiVec system indicated that tardigrade-specific genes are expressed specifically in different tissues, contrary to the previous expectations that these proteins are expressed and synergistically function within a single cell. CAHS genes are mainly expressed in epidermal cells, and the proteins are localized in the cytosol, while SAHS proteins are expressed exclusively in storage cells, which are tardigrade-specific free-floating cells in the body cavity. Therefore, the TardiVec system enables live imaging and transient expression in tardigrades and is especially useful in examining the details and dynamics of anhydrobiosis.

## Materials and Methods

### Animals

The YOKOZUNA-1 strain of *R. varieornatus*, the Z151 strain of *Hypsibius exemplaris*, previously described as *H. dujardini*^*38*^, and the PL.014 strain of *Thulinius ruffoi* were used. These strains were maintained on water-layered agar plates by feeding algae *Chlorella vulgaris* (Chlorella Industry)^4,39^.

### Design of the vector system and preparation of the injection solution

TardiVec systems were designed to consist of a promoter, mEGFP, 3’UTR, selection marker, and a replication origin for amplification in *E. coli*. The promoter and 3’UTR regions correspond to 1 kbp DNA sequences located before and after an open reading frame, respectively. The expression marker is a monomeric enhanced green fluorescent protein (mEGFP) fused with the nuclear localization sequence (NLS; PKKKRKV), and thus it facilitates the observation of fluorescence signals by concentrating them in the nucleus. The selection marker was ampicillin. Promoter, mEGFP and 3’UTR fragments were amplified by PrimeSTAR Max DNA polymerase (Takara). The replication origin and selection marker were derived from a mammalian expression vector, pCAGGS. pCAGGS was digested by *Spe*I and *Bam*HI and assembled with the PCR products by an In-Fusion HD cloning system (Clontech). After transformation into DH5α, the amplified plasmids were extracted by a Plasmid Maxi Kit (QIAGEN) to a concentration of approximately 2 μg/μL. The resulting pellets were eluted with pure water.

### Introduction of plasmids to tardigrades via microinjection

Glass capillaries (GD-1, NARISHIGE) were pulled by a puller (PC-100, NARISHIGE); the temperatures were set at 66.2 and 62.0°C. Our microinjection system consists of an inverted microscope (AXIO Vert. A1, Zeiss) equipped with an injector (IM-31, NARISHIGE) and manipulators (MMN-1 and MHW-103, NARISHIGE). Tardigrades were mounted as described previously by Tenlen et al. for RNAi experiments without anesthesia (see Figure 1C)^40,41^. After the microinjections, individuals were collected and transferred to a cuvette (CUY505P5, NEPA GENE) for electroporation using a super electroporator (NEPA21 type 2, NEPA GENE). The poring pulse was emitted twice at 250 V for 5 msec with a 50 msec interval, and the transfer pulse was emitted five times at 30 V for 50 msec with a 50 msec interval. Tardigrades were maintained with algal food on an agar plate until observation at 17°C.

**Figure 1.**
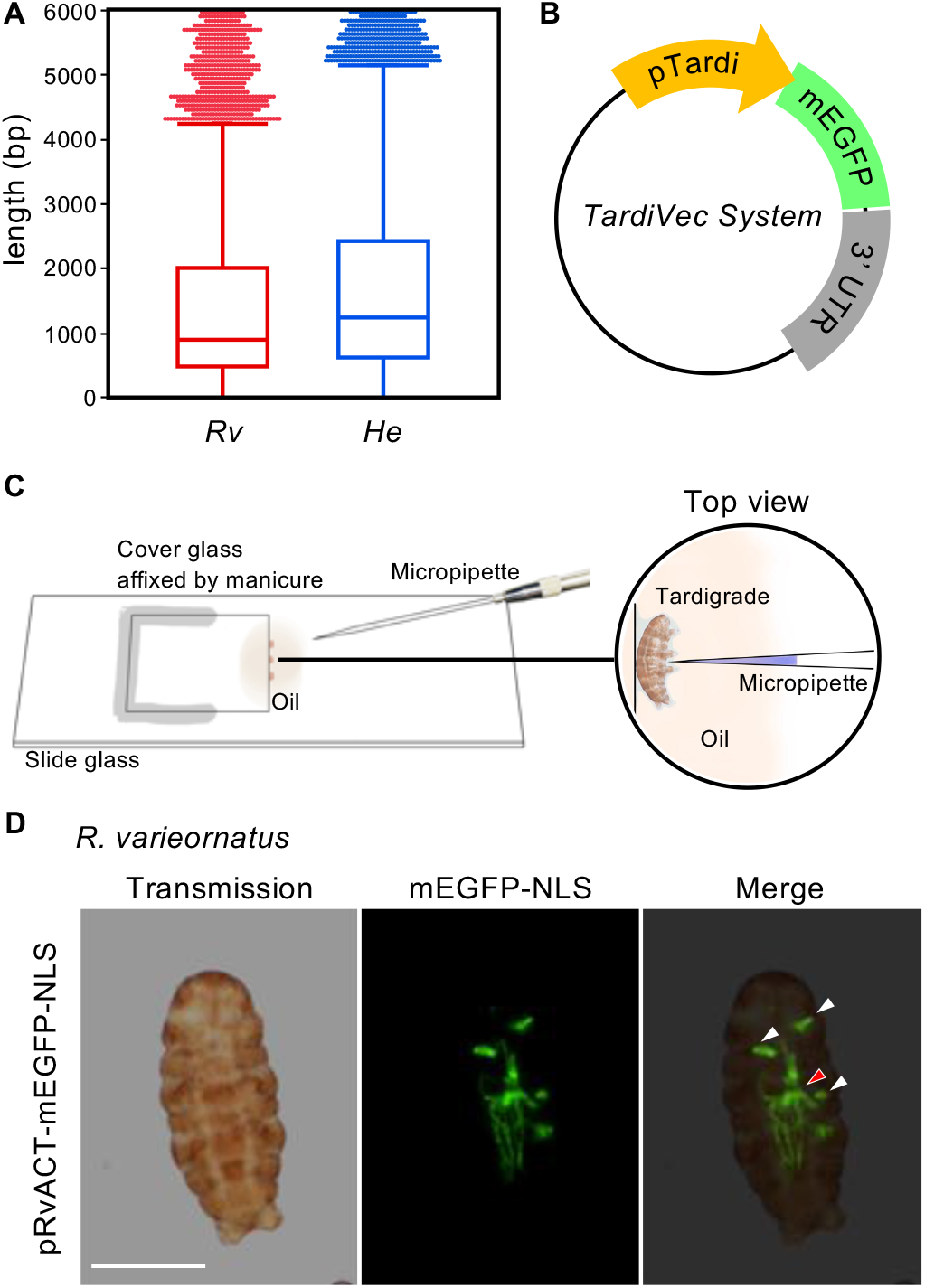
Tardigrade vector system enables *in vivo* expression in tardigrades. (A) Length of intergenic region in the genomes of *R. varieornatus* (*Rv*) and *H. exemplaris* (*He*). (B) Tardigrade vector named TardiVec. TardiVec consists of a promoter and terminator that originate from the genome of tardigrades. (C) Microinjection into tardigrades. To perform microinjection, a thin cover glass is affixed on the slide glass, and tardigrades are soaked in halocarbon oil with a small amount of water for fixation. (D) Tardigrade, *R. varieornatus*, expressing mEGFP-NLS under the actin promoter of *R. varieornatus*, pRvACT. White arrowheads indicate the mEGFP signal in muscle cells, and a red arrowhead indicates the signal in a neuron. Scale bars: 100 μm.

### Observation of fluorescence in tardigrades

Tardigrades were sandwiched between cover glasses for observation. Polystyrene particles were sometimes used to maintain the distance between the glasses. The bright field and fluorescence images were obtained using an IX73 inverted microscope with a DP74 camera adapter (Evident (Olympus)). LUCPLFLN 20X PH and LUCPLFLN 40X PH lenses (Evident (Olympus)) were used. For observation of GCaMP6s and subcellular localization, a TCS SP8 confocal laser scanning microscope (Leica Microsystems) equipped with HC PL APO CS2 20x/0.75, HC PL APO CS2 63x/1.20, and HC PL APO CS2 100x/1.40 objectives (Leica Microsystems) was used. Images were analyzed with ImageJ2 and Fiji.

### Detection of plasmid DNA from transfected tardigrade samples

Tardigrades transfected with pRvACT-mEGFP-NLS were kept for 10 days, and 10 individuals were collected at each time point as samples. Tardigrade samples were stored at -80°C until extraction treatment. An RNeasy mini kit (Qiagen) was used for extraction of the total nucleic acid. Amplification of target fragments was performed by PrimeSTAR Max DNA Polymerase (Takara).

### RNA sequencing

Storage cells were obtained by dissecting tardigrades in 0.1x PBS under a stereomicroscope with a sterile knife and quickly aspirating the cells that were released. Collected cells, as well as active individuals as controls for the whole body, were used to prepare RNA-Seq libraries using the SMART-Seq v.4 Kit (Clontech) as previously described^42^. Single-end sequencing was performed with a NextSeq 500 with a High Output Mode 75 Cycles Kit (Illumina). Expression levels were computed as transcripts per million (TPM) using kallisto software (v.0.46.1)^43^. The RNA-Seq data as well as computed abundances for each gene obtained were deposited into NCBI GEO under the accession ID GSE212632.

## Results

### Establishment of an *in vivo* expression system based on a plasmid vector with a tardigrade-specific promoter

To establish an *in vivo* expression system in tardigrades, we first designed DNA plasmids based on the expression system for housekeeping genes that are highly expressed in the transcriptome of *R. varieornatus*: *actin, ubiquitin*, and *tubulin*. The genome size of *R. varieornatus* is relatively small, 56 Mbp, and the median length of intergenic regions is less than 1 kbp (Figure 1A)^19,44^. Therefore, we extracted the 1 kbp sequence upstream and downstream of each target gene as the potential promoter and 3’UTR regions to activate and terminate transcription. The expression marker used was an mEGFP with an NLS sequence tag at the C-terminus (Figure 1B), so it was expected that concentrated fluorescence would be observed in the nucleus even if the expression level was quite low. As a result of the introduction of the plasmid vector to tardigrades (Figure 1C), we observed GFP fluorescence in tardigrades (Figure 1D and S1A). Fluorescence in most cells was observed to spread throughout the cell, although it was most intensive in the nucleus; it is thought that this is due to the overflow from the nucleus. Injection of the plasmid solution was insufficient for successful *in vivo* expression, and electroporation or transfection reagents were necessary. The survival rate after microinjection was almost 100%, but some individuals, approximately 0-20%, died after electroporation, presumably depending on the amount of injected solution. In more than 80% of the tardigrades treated with the TardiVec system, GFP fluorescence was observed after at least 24 hours (Table 1). We have so far observed GFP fluorescence in the muscle, epidermis, neuron, storage cells, and ovary, and the fluorescence in muscle cells was more intense than that in other tissues. This is probably because of the difference in the efficiency of introduction, as well as the promoter activity in each tissue. The variation of expression among individuals possibly depends on the differing membrane damage during electroporation, where the amount of DNA injected affects electrical resistance of tardigrade individuals and the corresponding membrane damage. Injection of low-concentration plasmid solution had a low success rate, but the high-concentration mixture of the two plasmids showed equal expression even though each concentration was halved. Transfection reagents were functional in tardigrade cells, but the toxicity was not negligible, and there was a tendency for expression to occur only in the periphery of the injection site. The plasmid DNA was detected in the tardigrades that were kept for 10 days after introduction, although they were degraded gradually (Figure 2A). GFP fluorescence was observed and remained strong in the individuals on the 10th day: intensity was comparable to that at 48 h and 72 h after introduction (Figure 2B). When the mEGFP inserted downstream of the actin promoter was replaced with the calcium indicator GCaMP6s, we observed blinking of fluorescence in the muscle cells and neurons, i.e., calcium oscillation associated with muscle contraction and neural activity (Figure 2C and Movie 1). The genetically encoded Ca^2+^ indicator can also function in tardigrades, and therefore, their *in vivo* calcium concentration is at a comparable level with other organisms (the Kd for Ca^2+^ of GCaMP6s is approximately 140 nM)^45^. We also observed that the GCaMP signal increased during dehydration, suggesting that the intracellular calcium concentration increased (Movie 2). To the best of our knowledge, this is the first direct observation of intracellular molecular changes in anhydrobiotic animals. Notably, GCaMP consists of circularly permuted GFP, with calcium-binding protein calmodulin at the C-terminus and the calmodulin interacting M13 peptide at the N-terminus; therefore, its functional blinking indicates that the transcription and translation of the inserted gene with a length of 1,251 bp was completely continuous from the N-terminal domain to the C-terminal domain in the TardiVec system.

**Table 1.** Survival rate and transfection rate upon introduction of each vector.

| Species | Vector | Concentration [$\mu\text{g}/\mu\text{L}$] | Survival rate | Transfection rate |
| --- | --- | --- | --- | --- |
| *R. varieornatus* | pRvACT-mEGFP-NLS | 1.9 | 96% (22/23) | 82% (18/22) |
|  | pRvUBC-mEGFP-NLS | 4.5 | 97% (30/31) | 97% (29/30) |
|  | pRvTUB-mEGFP-NLS | 2.6 | 100% (32/32) | 88% (28/32) |
|  | pRvCAHS3-mEGFP-NLS | 2.0 | 83% (25/30) | 83% (15/18)* |
|  | pRvSAHS1-mEGFP-NLS | 4.1 | 64% (23/36) | 91% (21/23) |
|  | pHeACT-mEGFP-NLS | 3.5 | 83% (24/29) | 100% (24/24) |
|  | pHeEF1a-mEGFP-NLS | 3.5 | 83% (25/30) | 80% (20/25) |
|  |  |  | Average 87% | Average 89% |
| *H. exemplaris* | pHeACT-mEGFP-NLS | 3.5 | 80% (28/35) | 61% (17/28) |
|  | pHeEF1a-mEGFP-NLS | 3.5 | 48% (12/25) | 67% (8/12) |
|  | pRvCAHS3-mEGFP-NLS | 2.0 | 70% (16/23) | 56% (9/16) |
|  | pRvSAHS1-mEGFP-NLS | 4.1 | 67% (16/24) | 100% (16/16) |
|  |  |  | Average 66% | Average 71% |

\*7 samples were lost just before the expression check.

**Figure 2.**
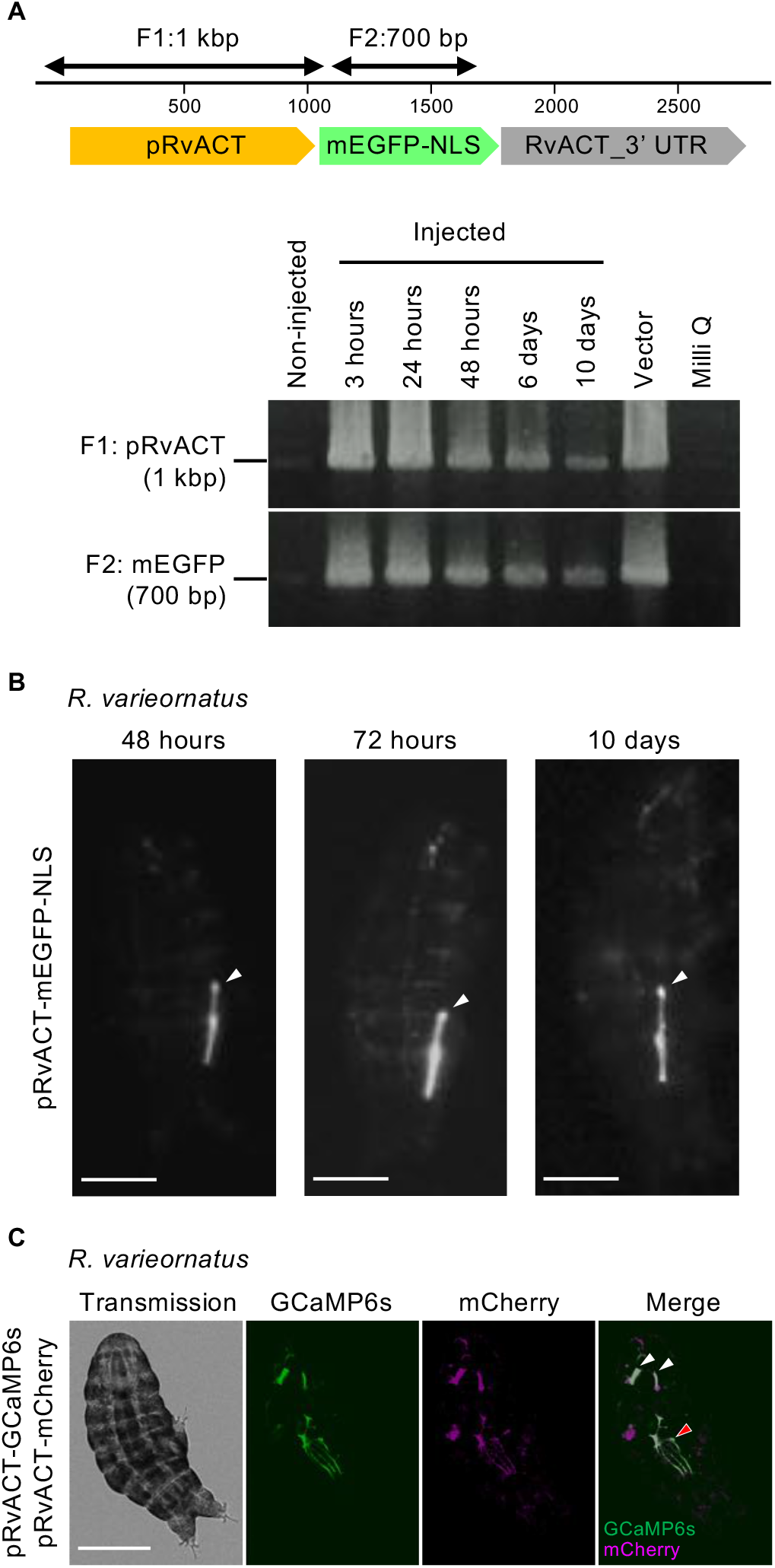
GFP fluorescence and plasmid DNA were retained for 10 days after introduction and GCaMP functioned in tardigrades. (A) Detection of plasmid DNA extracted at each time point from transfected tardigrades. The upper panel shows bands obtained from PCR targeting the pRvACT region (1 kbp), and the lower panel shows the mEGFP region (700 bp). Their locations in the pRvACT-mEGFP-NLS vector are represented in the right panel. (B) Time series images of the same tardigrades that were injected with pRvACT-mEGFP-NLS at 48 h, 72 h, and 10 days after injection. White arrowheads indicate the same muscle cell. Scale bars: 100 μm. (C) Tardigrades expressing GCaMP6s and mCherry with pRvACT. White arrowheads indicate the GCaMP signal in muscle cells, and a red arrowhead indicates the signal in a neuron. Scale bars: 100 μm.

Similar to *R. varieornatus*, we developed other DNA vectors based on the genome and transcriptome data of *H. exemplaris*, which is another anhydrobiotic tardigrade widely studied for development due to its transparency^39^, another suitable characteristic for fluorescence observation. These two species are closely related, and both belong to the superfamily Hypsibioidea, but there is a fundamental difference in the mode of anhydrobiotic entry; *R. varieornatus* constitutively expresses anhydrobiosis-related proteins to cope with rapid desiccation, whereas *H. exemplaris* slowly prepares and expresses these proteins during a period called “preconditioning” upon the detection of water loss^46^. Comparison between these two species is therefore important to understand the assembly of anhydrobiotic machinery and the evolution of anhydrobiosis. Although *H. exemplaris* has approximately twice the genome size of *R. varieornatus*, the median length of the intergenic region is only slightly longer at approximately 1,200 bp (Figure 1A)^44^; therefore, we adopted the same design as *R. varieornatus* in the *in vivo* expression of *H. exemplaris*. Actin and ef1α, which are highly expressed in *H. exemplaris*, were initially selected for the DNA vector. These promoters showed high expression of mEGFP ubiquitously in *H. exemplaris* (Figure 3A and S1B). Interestingly, we observed that these vectors also functioned interchangeably in *R. varieornatus*, and *vice versa*, where the vectors of *R. varieornatus* also functioned seamlessly in *H. exemplaris*, even though the identity rate of the actin promoter between these two species is only approximately 50% (Figure 3B and S1B).

**Figure 3.**
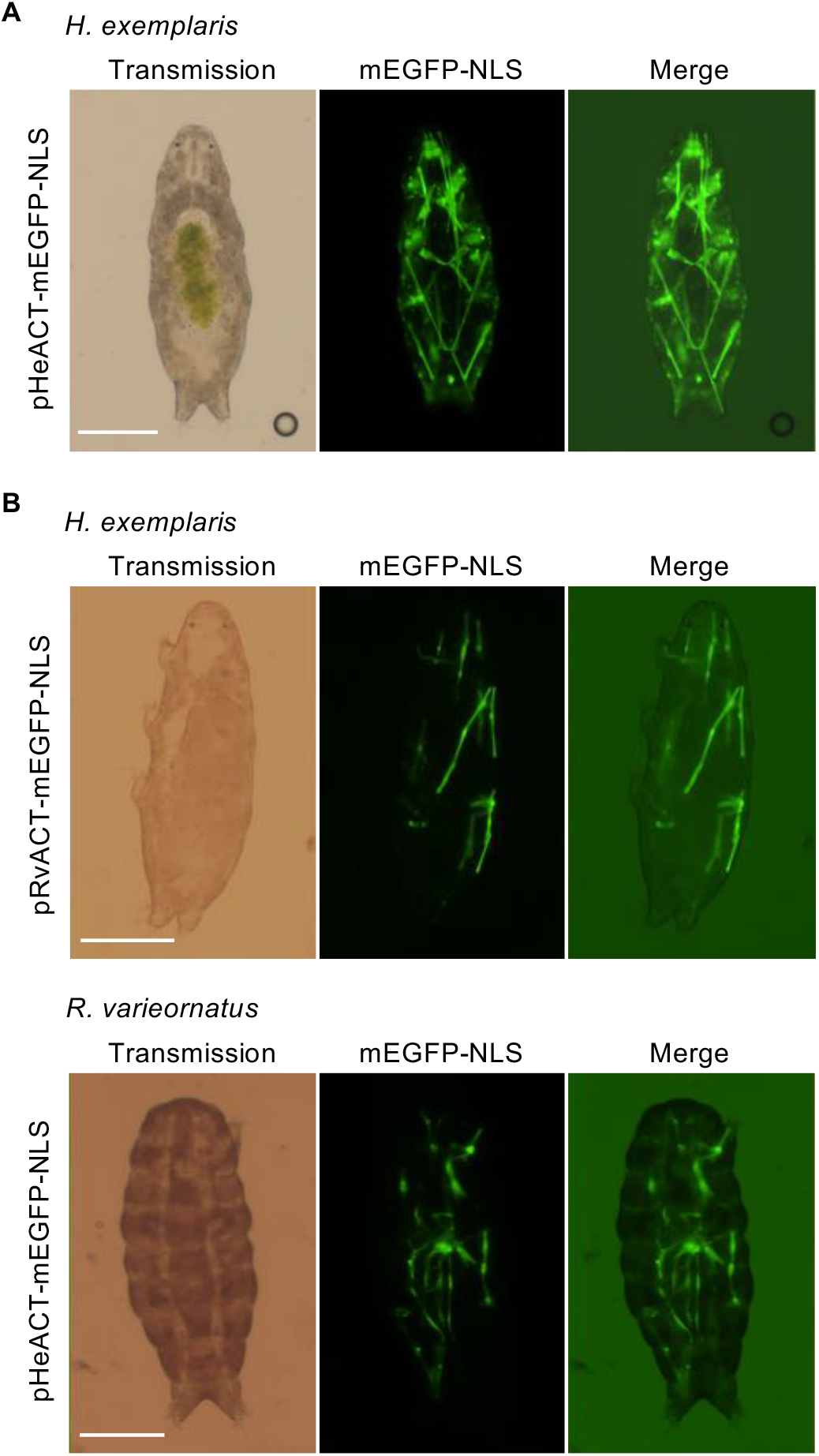
TardiVec systems are interchangeable within the same family. (A) Another anhydrobiotic tardigrade, *H. exemplaris*, expressing mEGFP-NLS under the actin promoter that originates from the genome of *H. exemplaris*. (B) *R. varieornatus* and *H. exemplaris* expressing mEGFP-NLS by interchange introduction of TardiVec. Scale bars: 100 μm. The merged images were obtained by adding transmission light to observe tardigrade body under the condition of observing GFP signal.

### Tissue-specific expression patterns of anhydrobiosis-related genes are conserved in anhydrobiotic tardigrades

Several tardigrade-specific genes have been discovered and then named based on their subcellular localization in human cultured cells, e.g., cytoplasm-localized CAHS1-3 and secretory SAHS1. They are IUPs highly expressed in tardigrades; therefore, they have been assumed to act as a “molecular shield” that has been hypothesized to be the main protective mode of anhydrobiotic protectants such as LEA proteins^47,48^. These proteins were previously assumed to function in concert; it is supposed that CAHS proteins protect the cytoplasmic components, while SAHS proteins protect the extracellular components, in coordination with other subcellular protectant proteins, MAHS and LEAM in mitochondria, and Dsup in the nucleus, altogether protecting all parts of the cells during anhydrobiosis. To test this “single cell hypothesis”, we developed vectors with promoters of *CAHS3* (pRvCAHS3), which is most highly expressed among *CAHS1-3*, and *SAHS1* (pRvSAHS1). As a result, tardigrades transfected with pRvCAHS3 plasmids showed intensive GFP signals in the epidermal tissue (Figure 4A). Even more unexpectedly, tardigrades injected with the pRvSAHS1 plasmid showed intensive expression most exclusively in the storage cells (Figure 4B). Most cells expressing the mEGFP-NLS protein under the RvSAHS1 promoter leaked out of the body when pressed, confirming that these are free-floating cells (Movie 3). The CAHS gene family consists of the *CAHS1/2* and *CAHS3* subfamilies; to examine the expression pattern of another *CAHS* gene subfamily, *CAHS1/2*, plasmid vectors of *CAHS1* were introduced into *R. varieornatus*. As a result, pRvCAHS1 showed the same expression pattern as pRvCAHS3 in epidermal cells (Figure S2A), suggesting that *CAHS* subfamilies possibly work together in the same cells rather than in different tissues. Tissue-specific expression of *SAHS* in storage cells was further confirmed, and this expression was validated directly through RNA sequencing of these storage cells because these cells could be collected by incision. The transcripts of the *SAHS* gene family were detected in much higher abundance compared to the whole body (Figure 4C), and the fold change (x5-20) roughly corresponded to the ratio of the number of cells in the storage cells to the whole body^49^.

**Figure 4.**
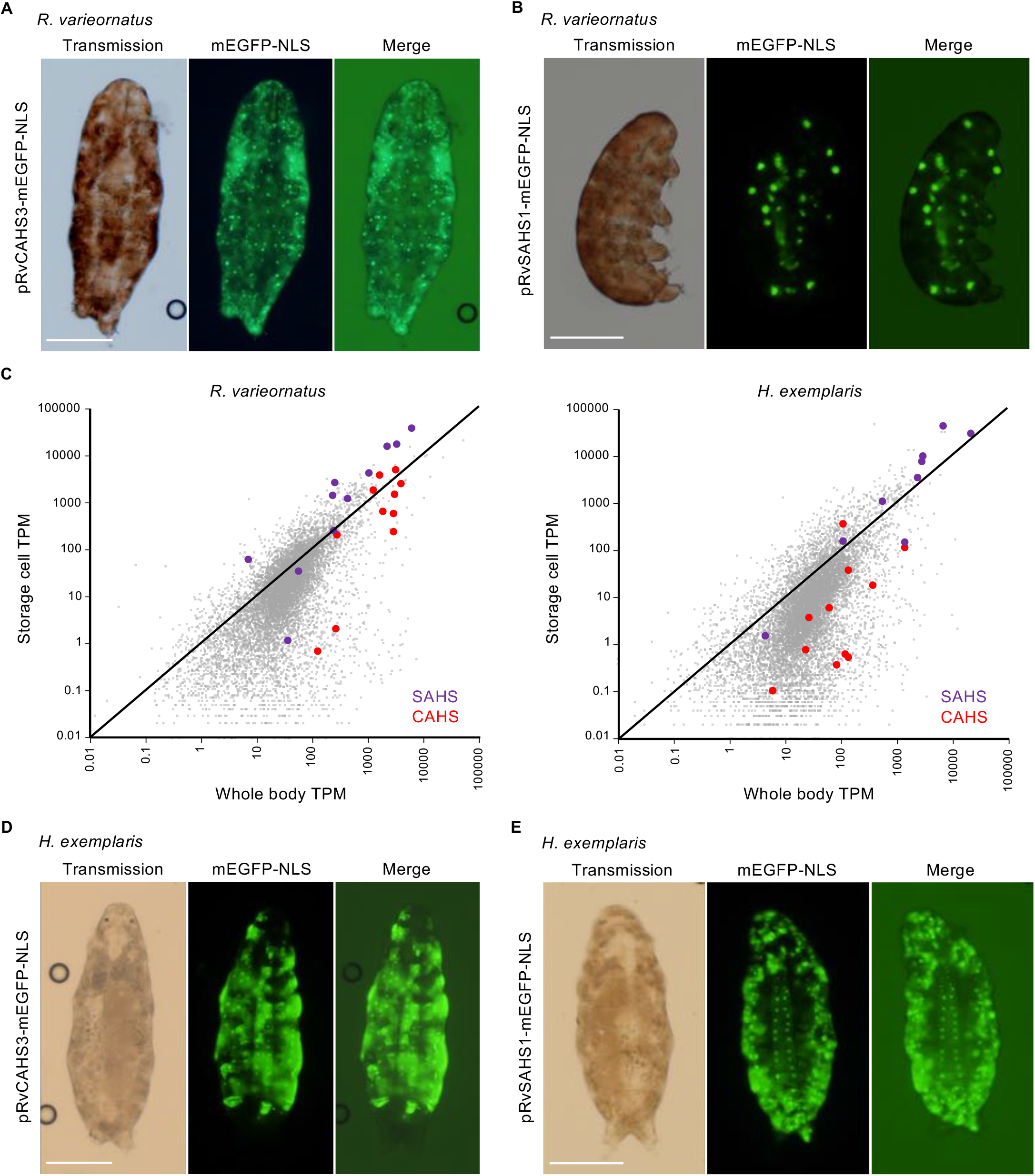
The tissue-specific expression of the tardigrade-specific genes *CAHS* and *SAHS* in *R. varieornatus* and *H. exemplaris*. (A) *R. varieornatus* expressing mEGFP under pRvCHAS3. (B) *R. varieornatus* expressing mEGFP-NLS under pRvSAHS1. (C) Comparison of transcripts of *CAHS* (red circle) and *SAHS* (purple circle) gene families between storage cells and the whole body of *R. varieornatus* and *H. exemplaris*. (D) *H. exemplaris* expressing mEGFP under pRvCAHS3. (E) *H. exemplaris* expressing mEGFP under pRvSAHS1. Scale bars: 100 μm. The merged images were obtained by adding transmission light to observe tardigrade body under the condition of observing GFP signal.

Surprisingly, these two plasmids also showed similar intense expression patterns in *H. exemplaris*, contrary to the premise that *H. exemplaris* requires a stimulation and preconditioning period to induce the expression of anhydrobiosis-related genes (Figure 4D, E, and Movie 3). Furthermore, the aquatic tardigrade *T. ruffoi* also expressed the pRvCAHS3 and pRvSAHS1 plasmids with similar tissue specificity (Figure S3). *T. ruffoi* belongs to the order Parachela with *R. varieornatus* and *H. exemplaris* but is highly sensitive to desiccation, and therefore was expected to lack the capacity to express *CAHS* and *SAHS*. Moreover, it was found that pHeCAHS (BV898_02951, CAHS 86272), which is composed of the 5’ and 3’ regions of the most highly expressed *CAHS* gene in the anhydrobiotic state of *H. exemplaris*, functions not only in *H. exemplaris* but also in *R. varieornatus* (Figure S2B). Taken together, these results indicate that the 1 kbp sequence of the 5’ region of anhydrobiosis-related genes contains the conserved elements required for its transcription and tissue specificity, but not for the regulation of transcription, at least in terms of anhydrobiotic response or suppression thereof.

### Subcellular localization and dynamics of anhydrobiotic proteins in tardigrade cells

Next, to confirm the subcellular localization and dynamics of CAHS and SAHS proteins in actual tardigrade cells, we introduced plasmids in which mEGFP-NLS was replaced with intact proteins fused with mEGFP, such as CAHS3-mEGFP or SAHS1-mEGFP, into pRvCAHS3 or pRvSAHS1 vectors. The CAHS3-mEGFP is localized in the cytoplasm, and fewer signals were observed in the nucleus, which is the same outcome as that reported in human cultured HEK293T cells, while the mEGFP-only control localized throughout the cytosol and in the nucleus (Figure 5A). To investigate the dynamics of these anhydrobiotic proteins when entering anhydrobiosis, we next observed tardigrades with pRvCAHS3-CAHS3-mEGFP introduced under desiccation. As a control, we prepared a vector to express only mEGFP and mCherry. Contrary to the report in cultured cells^27^, we did not observe filament formation of CAHS3-mEGFP in tardigrade cells (Movie 4). Since dehydration of the specimen made the fluorescence observation challenging due to cell compression, we then tested osmotic stress by soaking the tardigrades in 0.1 M NaCl. Under this osmotic stress condition, CAHS3-mEGFP seemed to accumulate along membranes, sometimes forming a fiber-like assembly, while mCherry coexpressed in the same cells remained uniform (Figure S4), and mEGFP also did not show any distinct change. This result thus suggests that CAHS3-mEGFP is possibly associated with the membrane in achieving anhydrobiosis. In addition, CAHS3-mEGFP and the mEGFP control showed LLPS-like behavior during observation (Movie 5), suggesting that the intracellular condition of tardigrades might be quite different from that of other experimental model organisms.

**Figure 5.**
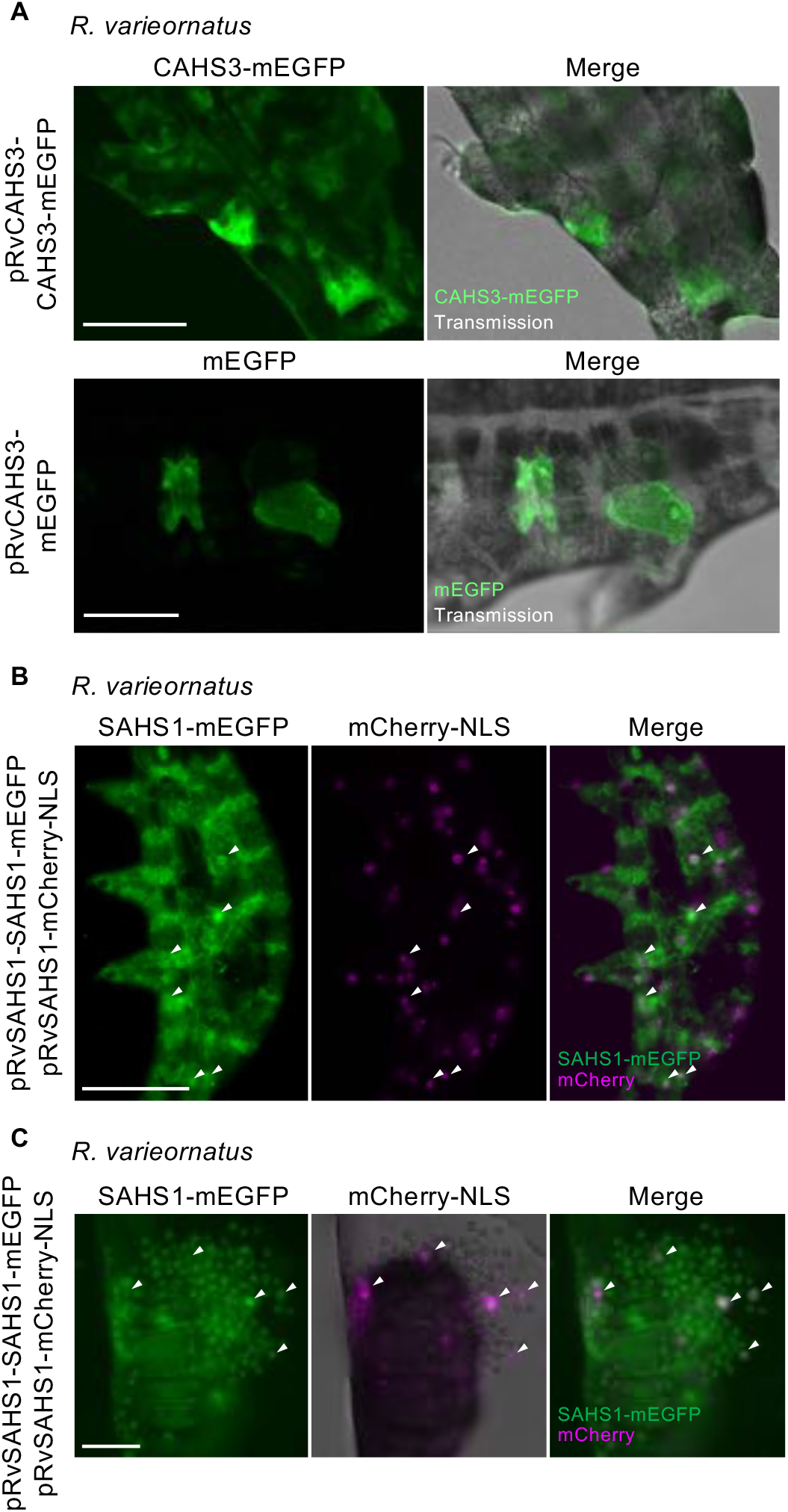
Subcellular localization of tardigrade-specific proteins fused with mEGFP in tardigrade cells. (A) Overexpression of CAHS3-mEGFP and mEGFP-only in *R. varieornatus*. (B) Overexpression of SAHS1-mEGFP and mCherry-NLS by cotransfection in *R. varieornatus*. (C) Observation of a punctured tardigrade expressing SAHS1-mEGFP and mCherry-NLS. White arrowheads indicate cells in which mEGFP and mCherry signals were merged. Scale bars: 50 μm.

On the other hand, SAHS1-mEGFPs were observed in the vesicle-like structure in the storage cells, while in some individuals, GFP signals were observed to spread throughout the body cavity (Figure 5B and S5A). This pattern is similar to the previous heterologous expression observed in HEK293T cells, in which weak signals in secretory pathways such as the endoplasmic reticulum or the Golgi apparatus were observed and most SAHS1-mEGFP was detected from the culture media. The fact that most SAHS1-mEGFPs were retained in the storage cells in tardigrades suggests differences in the modes of secretion and storage regulation between tardigrades and human cells, especially since the cell type is unique to tardigrades. Localization of SAHS1-mEGFP in the storage cell was again confirmed by leaking the cell contents out by puncture (Figure 5C). These results demonstrated that the CAHS proteins localized in the cytoplasm of the epidermal tissue, and the SAHS proteins were stored in vesicles of the body cavity cells in tardigrades. Moreover, the subcellular localization was also conserved in *H. exemplaris* (Figure S5B and C).

## Discussion

### Inception of live imaging in anhydrobiotic tardigrades

Tardigrada is one of the phyla in Ecdysozoa; it is still controversial whether Tardigrada should be included in Panarthropoda due to controversial molecular evidence suggesting similarity with Cycloneuralia ^44^. Nevertheless, there are model organisms with modern genetic toolkits around the phylum Tardigrada in the ecdysozoan tree of life, such as *C. elegans* and *D. melaogaster*. Such an experimental model species was lacking in Tardigrada, even though the phenomenon of anhydrobiosis in tardigrades has been enthralling researchers for decades. Following the establishment of the stable culture system, complete genome sequences have been published for *R. varieornatus* and *H. exemplaris* over the past decade, and as a result, several tardigrade-specific genes have been discovered to contribute to anhydrobiotic machinery. Our TardiVec system achieves prolonged expression in various tissues across different orders within Tardigrada, with the capability to cotransfect multiple vectors at the same time. It is therefore expected to contribute not only to the study of these anhydrobiosis-related proteins, but also to various studies of tardigrades, including development, physiology, and ethology. Tardigrades have thus become an emerging model species.

The TardiVec system easily enables the introduction of live imaging with tags (e.g., GFP) or indicators (e.g., GCaMP). Fluorescent proteins fused with target proteins possibly enable us to reveal not only the subcellular localization but also the protein dynamics during dehydration and rehydration without sample loss. For example, GCaMP will reveal neural activity and networks, building on the tardigrade anatomy that was revealed through IHC and electron microscopy imaging^29–36,50,51^.

Of note, the TardiVec system can work beyond the genus, the family, and the order, indicating that it can be readily applied to field-collected samples that have no precise genome data and adequate rearing systems. Indeed, we succeeded in the introduction of our vector plasmids that originated from *R. varieornatus* to anhydrobiotic tardigrades of another order, Apochela, *Milnesium inceptum*, which were collected from the field (Figure S6). The class Heterotardigrada, in which there is no isolated rearing system established yet for any of the species, has only a draft genome sequence of *Echiniscus testudo*^*52*^. The same vector system based on this species should also work across orders within heterotardigrades, opening up new frontiers in Tardigradology.

As shown in Figure 2, vector plasmids and GFP fluorescence were maintained for 10 days, suggesting that there is no active degradation mechanism by nucleases. Since cell division rarely occurs in tardigrades^53^, plasmids may not be lost by cell division as they are in cultured cells. Moreover, the half-life of GFP as a protein is approximately 26 hours^54^; therefore, it is reasonable to assume that the plasmid vector is functional up to 10 days after introduction. Although the length of the open reading frame is relatively short in *R. varieornatus*, the vector system could transcribe more than 1.6 kbp of product as reported here (CAHS3-mEGFP) and at least 2.5 kbp of preliminary product (data not shown). We conducted cotransfection using two distinct vectors (e.g., pRvCAHS3-CAHS3-mEGFP and pRvCAHS3-mCherry), and these two fluorescence signals were observed in almost entirely the same cells. This is possibly because plasmid DNA is imported selectively into cells with sufficient membrane damage by electroporation.

Recently, a CRISPR/Cas9 system functioning with gRNA and Cas9 proteins was reported in tardigrade somatic cells^55^. This system unfortunately did not seem to function in the germline, which is necessary to obtain a mutant strain by gene editing. Introduction of constructs to germlines is also a rare event in our method, but the TardiVec system will possibly provide another approach to deliver the gene-editing system to the germline, paving the way to produce gene-edited strains in tardigrades.

### Conservation and diversity of the expression mechanism of genes constituting anhydrobiotic machinery

Our present study showed that vectors derived from tardigrade-specific genes, i.e., *CAHS* and *SAHS*, successfully expressed inserted genes in *R. varieornatus, H. exemplaris*, and *T. ruffoi*. These results indicate that the 1 kb sequence upstream and downstream of each gene contains functional promoter and terminator sequences for constitutively expressing them with tissue specificity. The RvCAHS genes are constitutively expressed, while the expression of HeCAHS genes is induced only after dehydration cues; e.g., the fold change of HeCAHS is x20-x500 compared to the normal condition. Therefore, the pHeCAHS vector was expected to be nonfunctional under normal conditions of *H. exemplaris*, but such suppression was not observed; therefore, this result suggests that the regulation of the expression of these genes might be suppressed *in tarns* by repressor binding on distal regions and/or chromosomal regulation, partially suggested by the fact that these anhydrobiotic genes often colocalize within the genome in tandem. Moreover, this kind of global or meta-regulation of anhydrobiosis-related genes is in line with the fact that the shift from rapid anhydrobiosis to preconditioning or the loss of anhydrobiotic ability in Tardigrada seems to occur in multiple phylogenetic lineages and is a recurring adaptive strategy to cope with xeric, mesic, and aquatic environments^16^. It is obviously more energetically cost effective to only require the change in global regulation rather than evolving different transcription factor-promoter network rewiring for all related genes. Why is *CAHS* or *SAHS* expression conserved in *T. ruffoi*, which seemingly lost its anhydrobiotic capability? One possibility is another dormant state of tardigrades, which is known to be present in *T. ruffoi*, to form a cyst, an adaptive state to withstand unfavorable environmental conditions, although this corresponds with loss of anhydrobiotic ability^56^. The remaining promoter activity for anhydrobiosis-related genes in *T. ruffoi* allowed us to speculate regarding the possibility that the two dormant states of tardigrades, anhydrobiosis and encystment, are in fact somewhat related.

This “master switch” type of large-scale transcriptional regulation can be seen in the desiccation tolerance of plants. Plants can also adapt to water loss to some extent, and it has been revealed that the regulatory pathway induces resistance genes with complex crosstalk between transcription factors^57^. Overexpression of one of the transcription factors conferred drought resistance to plants by activating resistance genes^58^. A similar strategy could therefore be applicable to inducible anhydrobiotic tardigrades, such as *H. exemplaris*. Kondo et al. identified D942, an indirect activator of AMPK in mammals but not in tardigrades, as a candidate drug to regulate the expression of anhydrobiosis-related genes^59^.

### Tissue-specific expression pattern of tardigrade-specific proteins suggests systematic organismal anhydrobiosis machinery

Our study revealed that the expression of tardigrade-specific genes varied in specific tissues, although it had been assumed that all cells express the full set of anhydrobiotic genes. This single-cell hypothesis is based on the experimental results on human cultured cells; CAHS proteins localized in the cytosol and SAHS proteins were secreted into the medium^17,18^. As our results showed, these characteristics of subcellular localization in human cells are coincident with those in tardigrade cells, as was also reported for the subcellular localizations of RvLEAM and Dsup proteins by IHC of tardigrade embryos. The tissue-specific expression of these genes was revealed for the first time by whole-body observation with the TardiVec system, suggesting a more complex and systematic organismal anhydrobiosis machinery with a specialized set of proteins functioning in each of the specific tissue types. Of course, this does not rule out the fact that some secretory proteins are produced in a specific cell type but are spread throughout the body. In sleeping chironomids, trehalose, the main protectant molecule, is synthesized in the fat body, secreted to body fluid, and eventually spread throughout the whole body. Similarly, it is reasonable to assume that SAHS proteins are synthesized in storage cells and are secreted into the body cavity. Moreover, it is interesting that in most tardigrade samples, SAHS proteins seemed to remain in storage cells in our observation, whereas in rare cases, SAHS proteins were also observed to be present in the body cavity (Figure S5A). This suggests a regulatory mechanism governing secretion from cells in addition to transcription and translation. Additionally, since not all cells are in contact with the body cavity, it is possible that other mechanisms exist to protect the outside of the cells.

Although the single-cell hypothesis is attractive for establishing technologies for dry preservation of cells or other biological materials, the actual anhydrobiotic mechanism of tardigrades appears to be more complex; CAHS proteins mainly function in the epidermis, and SAHS proteins are produced in storage cells and possibly transported to other cells via the body cavity. Tissue-specific expression of each anhydrobiotic gene allows us to speculate with respect to the existence of other candidate proteins for accomplishing anhydrobiosis in various cell types. Tardigrade genomes are found to code multiple IUPs, and dozens of IUPs are highly expressed in *R. varieornatus* and induced in *H. exemplaris*, similar to *CAHS* and *SAHS* genes^15^; it is reasonable to assume that they are candidates as protectants of other cell types. In addition, the tissue specificity forces us to reconsider the sufficient amounts of protectants for understanding anhydrobiosis correctly.

Expression of the *CAHS3* gene by the TardiVec system was most often observed in epidermal cells. Since there are more than 10 paralogs of *CAHS* genes, there is still a possibility that the other paralogs of the *CAHS* family are expressed in other tissue types, or that all CAHS proteins may have a specific role in epidermal cells. Fibril formation was not confirmed in the observation of tardigrades expressing CAHS3-mEGFP, although reversible fiber or aggregation formation of CAHS proteins has been reported in animal culture cell lines^27^. Since the variety of *CAHS* paralogs makes us speculate that CAHS proteins are used differently based on their characteristics, future studies should analyze in more detail the differences in *CAHS* subfamilies and their tissue specificity. In addition, difficulty was encountered in observing the fluorescence of dehydrated samples due to insufficient light penetration and possible change in refractive index, but nevertheless, fluorescence observation is possible in the process of dehydration and rehydration. It has also been observed that GFP expressed in tardigrade cells exhibits LLPS-like dynamics during the dehydration process, suggesting that the condition of the cytoplasm might differ from that of other organisms. The fact that it became possible to observe the live cells of anhydrobiotic animals *in vivo* has enabled us to examine various possibilities to understand life without water.

To conclude, we developed a vector-based *in vivo* expression system consisting of tardigrade-specific promoters, named the TardiVec system. The TardiVec system enabled live imaging in several tardigrades in the class Eutardigrad, and the basic concept should be directly applicable for heterotardigrades as well. With this system, the dynamics of proteins and cells were revealed through observation using fluorescent tags and indicators in anhydrobiotic tardigrades. Live imaging would open new doors in the study of anhydrobiosis and make it possible to utilize tardigrades as a model system for anhydrobiology, morphology, physiology, and other various fields of biology.

## Acknowledgments

The authors are grateful to Esraa Hassan Ahmed Youssef for thoroughly conducting experiments associated with plasmid vectors. Naoko Ishii, Ayako Shirahata, Yuki Takai, and Takahiro Bino provided technical assistance. The *C. vulgaris* used to feed the tardigrades was provided courtesy of Chlorella Industry. This work is supported by KAKENHI Grant-in-Aid for Transformative Research Areas (A), Grant-in-Aid for Early-Career Scientists, and Grant-in-Aid for Challenging Research (Exploratory) from the Japan Society for the Promotion of Science (JSPS, grant Numbers 21H05279, 20K15781, and 22K19302), Joint Research by Exploratory Research Center on Life and Living Systems (ExCELLS program Nos. 19-208 and 19-501) and partly by research funds from the Yamagata Prefectural Government and Tsuruoka City, Japan.

## Author contributions

Conceptualization, S.T., and K.Ar.; Investigation, S.T. and K.Ar.; Writing – Original Draft, S.T.; Writing – Review & Editing, S.T., K.Ao., and K.Ar.; Funding Acquisition, S.T., and K.Ar.; Resources, S.T., K.Ao., and K.Ar.; Supervision, S.T. and K.Ar.

## Figure legends

Movie 1 | GCaMP6s and mCherry. The left panel is a merged bright field displaying GCaMP6s (green) and mCherry (red). The right panel shows merged GCaMP6s (green) and mCherry (magenta) (4x speed). GCaMP6s signal appear in a neuron at the point of 00:05 of the Movie 1.

Movie 2 | A tardigrade expressing GCaMP6s was dehydrated and rehydrated (16x speed). Water was added to rehydrate the tardigrade at 00:02.

Movie 3 | *R. varieornatus* and *H. exemplaris* expressing mEGFP-NLS under pRvSAHS1 (2x speed). In *R. varieornatus*, storage cells expressing mEGFP-NLS were ejected by pressure.

Movie 4 | Dehydrated tardigrades expressing (A) CAHS3-mEGFP and (B) mEGFP were rehydrated (16x speed).

Movie 5 | Dynamics of (A) CAHS3-mEGFP and (B) mEGFP in tardigrade cytosol (8x speed).

