## Supplementary material for "*in vivo* expression vector derived from anhydrobiotic tardigrade genome enables live imaging in Eutardigrada": Figure S

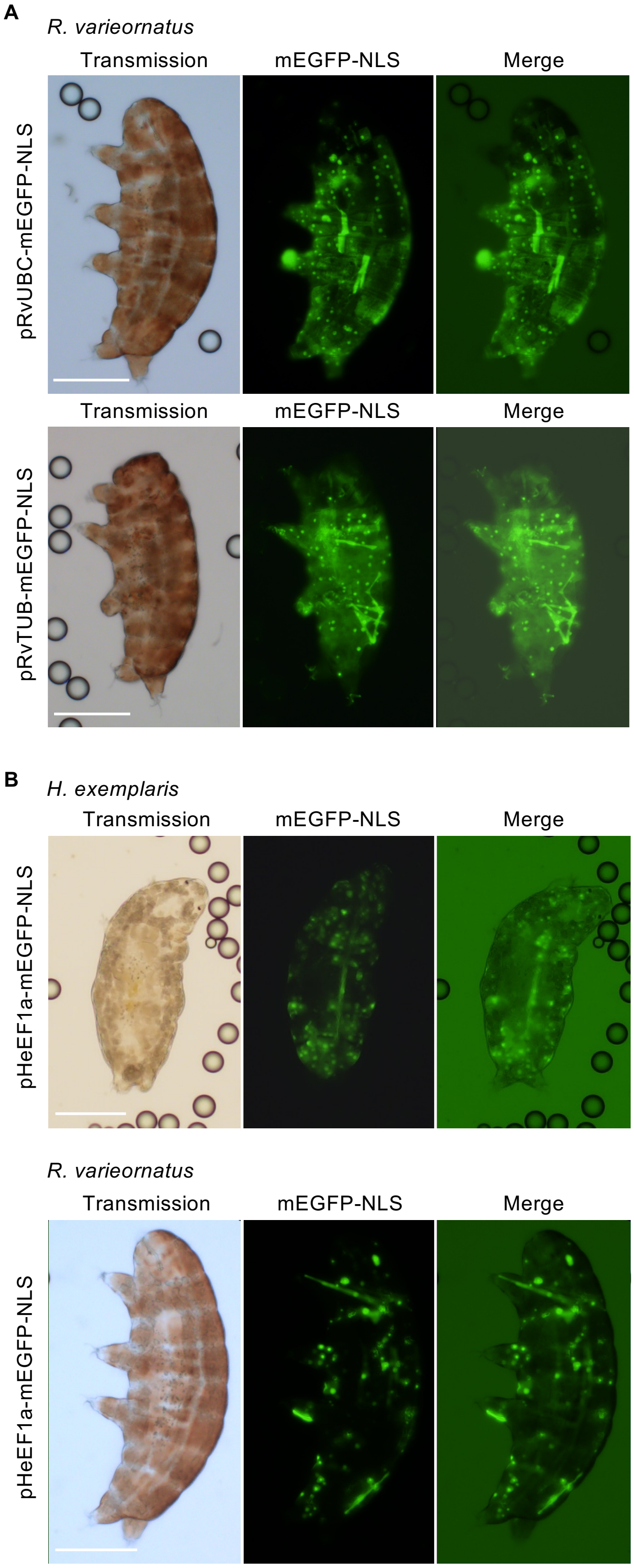


Figure S1 | (A) The promoters of the other housekeeping genes, *ubc* and *tubulin*, of *R. varieornatus* enabled mEGFP-NLS expression *in R. varieornatus*. Scale bars: 100 µm.

(B) Another *H. exemplaris* vector of the housekeeping gene *ef1α* enabled mEGFP-NLS expression in *H. exemplaris* and *R. varieornatus*. Scale bars: 100 µm.

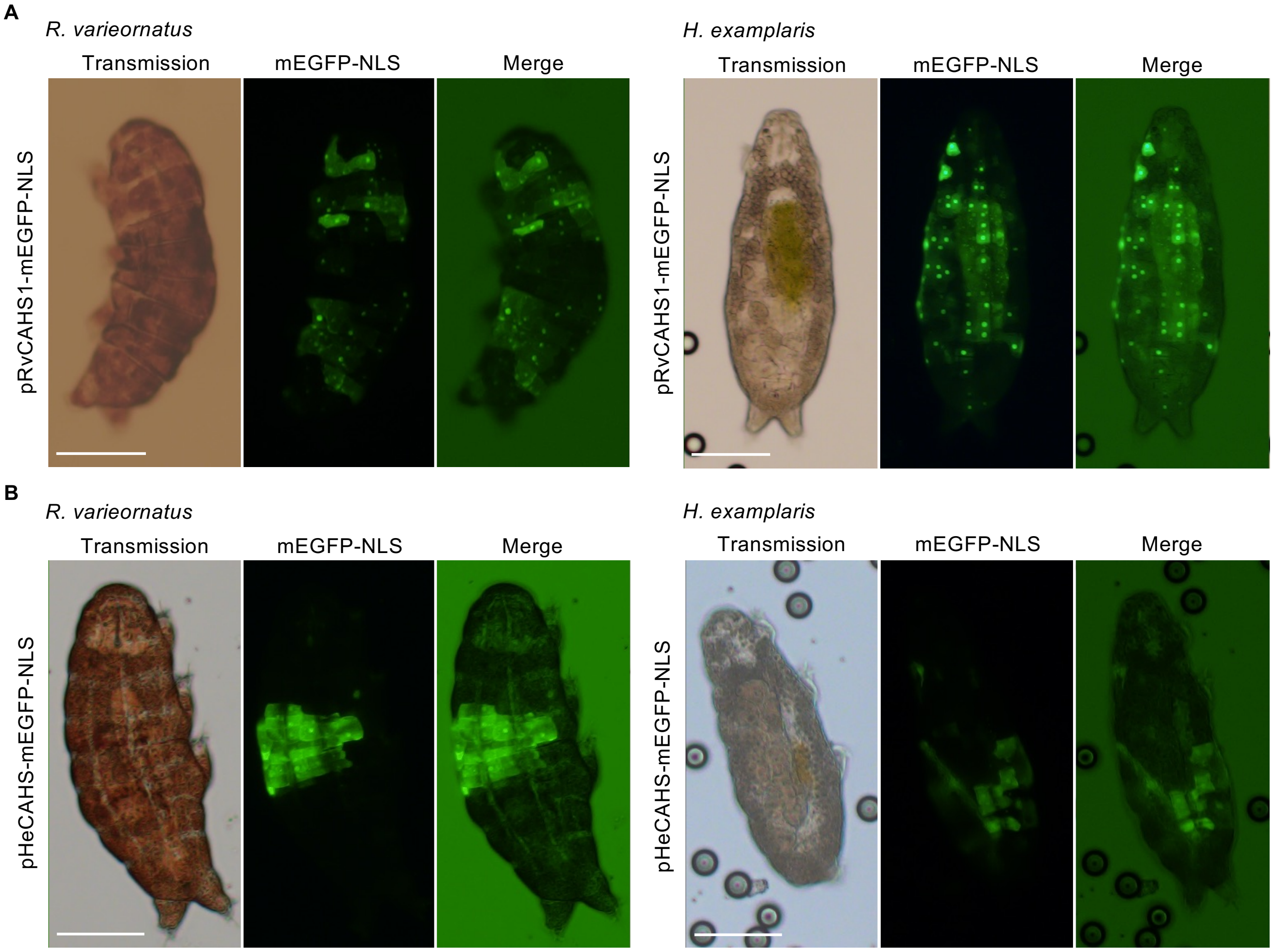


Figure S2 | Expression pattern of promoters of (A) *CAHS1* of *R. varieornatus* and (B) *CAHS* of *H. exemplaris* in *R. varieornatus* and *H. exemplaris*. Scale bars: 100 µm.

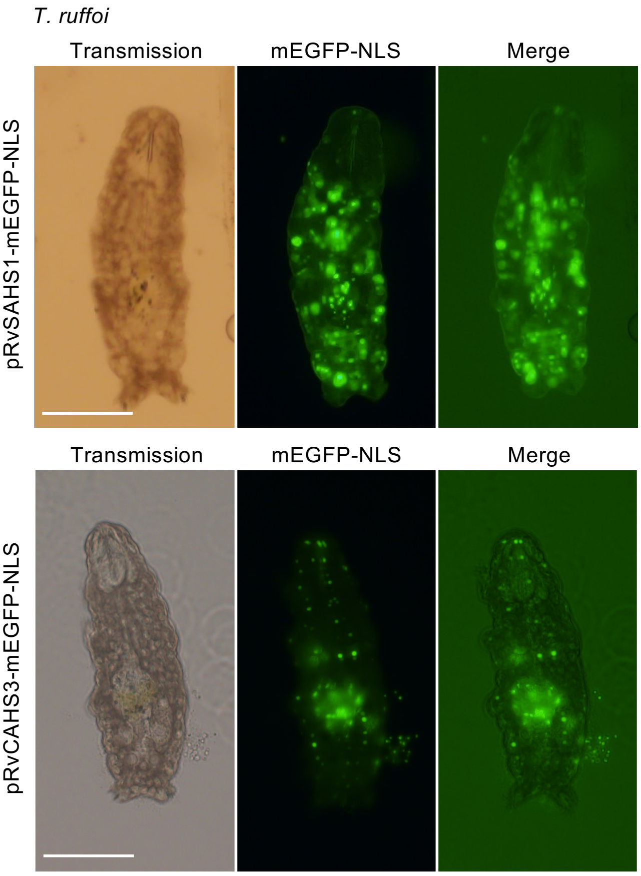


Figure S3 | Tardigrade-specific gene promoters function in non-anhydrobiotic tardigrades, *T. ruffoi*. Scale bars: 100 μm.

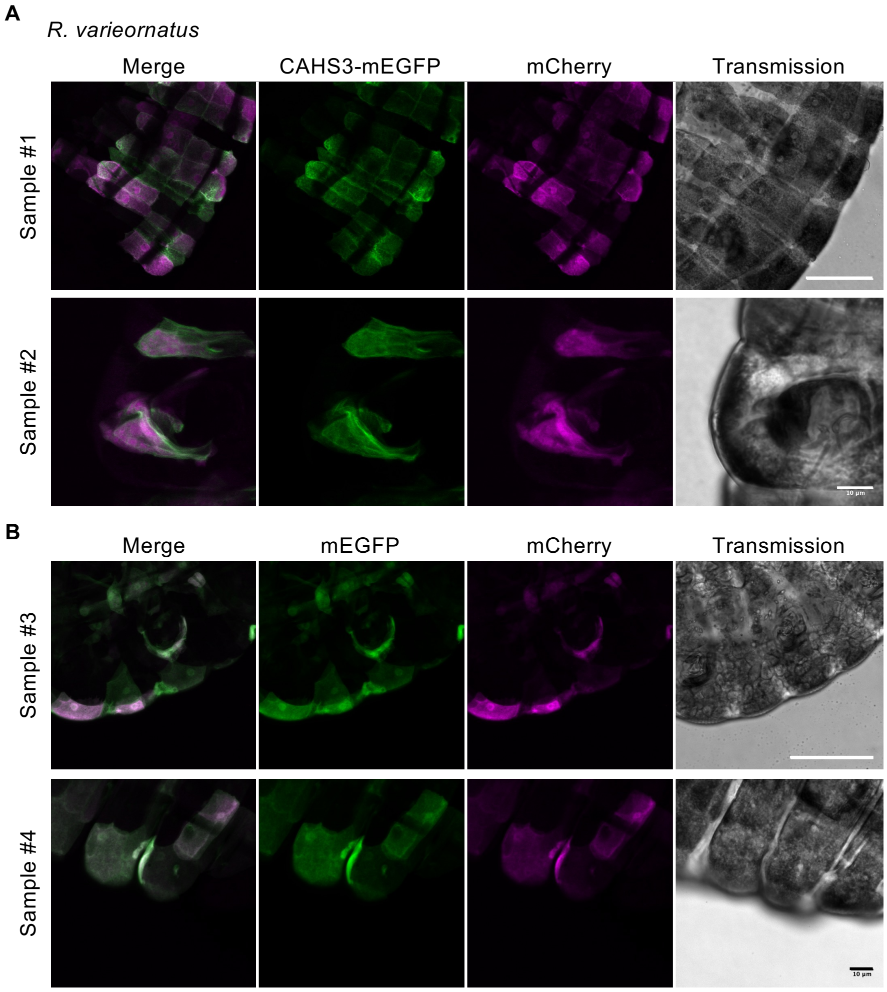


Figure S4 | Tardigrade in 0.1 M NaCl expressing (A) CAHS3-mEGFP and mCherry and (B) mEGFP-only and mCherry. Scale bars in sample #1 and #3: 50 μm, and in sample #2 and #4: 10 μm.

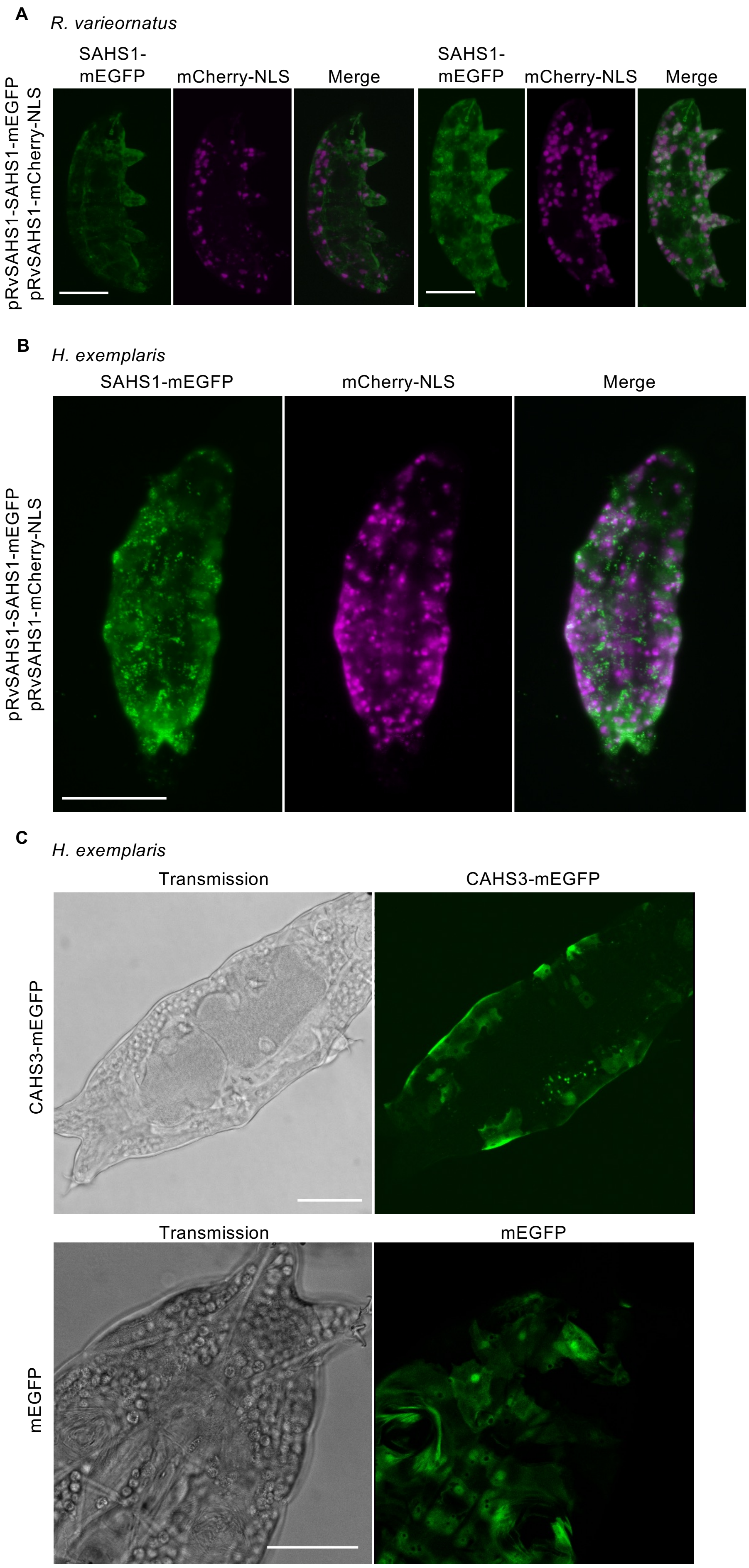


Figure S5 | Overexpression of (A) SAHS1-mEGFP in *R. varieornatus,* (B) SAHS1-mEGFP in *H. exemplaris*, and (C) CAHS3-mEGFP and mEGFP-only in *H. exemplaris*. Scale bars in A and B: 100 μm, and in C: 50 μm.

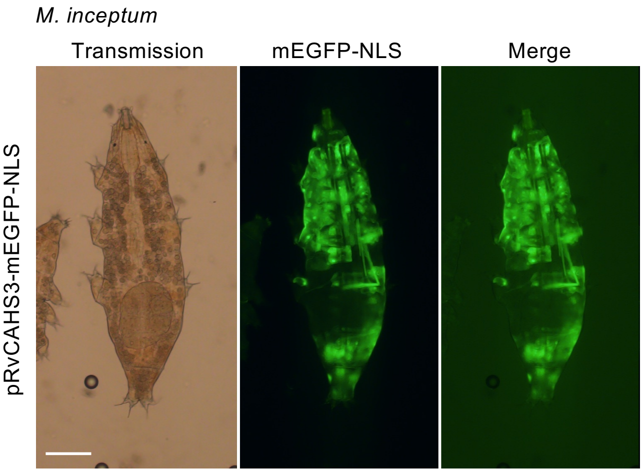


Figure S6 | The tardigrade-specific gene promoter, pRvCAHS3, functioned in Apochela, the other order of the class Eutardigrada. Scale bars: 100 μm.
